# A genomic baseline for long-term monitoring of economically important aquatic species in the Netherlands

**DOI:** 10.64898/2026.10.29.762745

**Authors:** Christopher D. Barratt, Sophie Valk, Elianne van der Valk, Annemieke Rattink, Wout Abbink

**Affiliations:** Centre for Genetic Resources the Netherlands, Wageningen; Animal Breeding and Genomics, Wageningen; Wageningen Marine Research, Wageningen

**Keywords:** anthropogenic pressure, aquaculture, climate change, food security, sustainability

## Abstract

Aquatic species of economic and ecological importance are increasingly threatened by environmental change, yet their genetic diversity and population structure remain poorly characterised, limiting effective conservation and management. To address this gap, we initiated a multi-species genomic monitoring programme in 2022, targeting key marine and freshwater taxa. A total of 2,152 individuals from 113 locations were sampled across six species: turbot (*Scophthalmus maximus*), zander (*Sander lucioperca*), European eel (*Anguilla anguilla*; both glass eel in the marine environment and adult eel in freshwater environments), blue mussel (*Mytilus edulis*), flat oyster (*Ostrea edulis*), and Pacific oyster (*Magallana gigas*). Using genotyping-by-sequencing (GBS) and SNP arrays, we generated genome-wide SNP datasets to assess population structure, genetic diversity, inbreeding, and relatedness. Among fish species, population structure was generally weak, consistent with high connectivity. Genetic diversity and relatedness varied among some sampling locations, highlighting the value of location-specific baselines for identifying patterns that warrant further investigation and may inform future conservation and management. In bivalves, genetic differentiation varied among species and sampling locations. Mussel and flat oyster showed differentiation among several sampled populations, including distinct cultured flat-oyster groups, whereas Pacific oyster showed comparatively weak structure in the analyses. Together, these results provide a genomic baseline for key aquatic species in the Netherlands and demonstrate the value of coordinated, multi-species genomic monitoring. The identification of genetically distinct and diverse populations offers direct guidance for conservation prioritisation, restoration strategies, and the sustainable management of aquatic genetic resources under ongoing environmental change, in the Netherlands but also applied in other national and transnational monitoring contexts.

## Introduction

Maintaining genetic diversity is a cornerstone of biodiversity conservation, providing the raw material for populations to adapt to environmental stressors such as climate change, disease, and habitat fragmentation (Dewoody et al., 2021; Hughes et al., 2008; Kardos & Luikart, 2021; Moritz, 2002). Beyond its evolutionary role, genetic diversity underpins key ecological processes, including productivity, nutrient cycling, and food-web stability. Empirical studies demonstrate that genetically diverse populations are more resilient to disturbance and better able to recover, thereby supporting long-term ecosystem function (Reusch et al., 2005; Reynolds et al., 2012). However, recent global analyses indicate that genetic diversity is declining across a wide range of taxa and is strongly associated with increased extinction risk, even in populations not currently classified as threatened (Shaw et al., 2025). Furthermore, genetic diversity is an essential component in safeguarding food production in general, to ensure sustainable breeding and resilience against climate change, pests and diseases, in crops (Renard & Tilman 2019), livestock species (Doeschl-Wilson et al. 2021) and aquatic species (Prunier et al. 2023). Safeguarding genetic diversity is therefore essential not only for species persistence but also for maintaining ecosystem integrity and the services upon which human societies depend (Formenti et al., 2022).

In aquatic systems, genetic diversity plays a central role in sustaining ecosystem health, supporting fisheries and aquaculture, and maintaining the ecological and socio-economic value of aquatic species. Aquatic genetic resources (AqGR) are thus critical for both biodiversity conservation and global food security. Recognising this, the Food and Agriculture Organization (FAO) adopted the Global Plan of Action for the Conservation, Sustainable Use and Development of Aquatic Genetic Resources for Food and Agriculture (FAO, 2015), which calls for the implementation of genetic monitoring programmes. Despite this, systematic genomic monitoring of aquatic species remains limited in most countries, constraining our ability to detect changes in genetic diversity and to implement timely, evidence-based management (Hoban et al., 2022, 2023; Laikre et al., 2020; Pearman et al., 2024; Thurfjell et al., 2022).

Recent European initiatives highlight the potential of genomic monitoring for aquatic management. For example, Sweden has begun integrating genetic data into fisheries assessments for species such as Atlantic herring (*Clupea harengus*) and cod (*Gadus morhua*), while Denmark is incorporating population genomics and eDNA approaches into stock identification and traceability frameworks. These efforts demonstrate the feasibility and value of genomic monitoring, but broader implementation across species and regions remains limited.

To address this gap, a multi-species genomic monitoring programme was initiated in 2022 targeting key marine and freshwater species: blue mussel (*Mytilus edulis*), flat oyster (*Ostrea edulis*), Pacific oyster (*Magallana gigas*), turbot (*Scophthalmus maximus*), zander (*Sander lucioperca*), and European eel (*Anguilla anguilla*). These species are of high economic and ecological importance, contributing to fisheries and aquaculture while playing key roles in ecosystem functioning. They are also subject to multiple anthropogenic pressures, including habitat degradation, overexploitation, pollution, and climate change, which may erode genetic diversity and adaptive potential. By encompassing a range of habitats and life histories, these species provide a representative framework for assessing aquatic genetic diversity across the Dutch landscape.

Here, we present the first genomic baseline for these species, based on samples collected across freshwater and marine systems in the Netherlands between 2022 and 2024. This baseline establishes a foundation for long-term genetic monitoring, enabling the detection of temporal trends and supporting proactive conservation, fisheries management, and restoration strategies. More broadly, this study demonstrates how coordinated, multi-species genomic monitoring can contribute to the conservation and sustainable use of aquatic genetic resources and provides a scalable framework for integrating genomics into biodiversity monitoring programmes.

## Materials and Methods

### Target species and sampling

Fish sampling was conducted in 2022 and 2024 from fishing vessels for turbot (*Scophthalmus maximus*), zander (*Sander lucioperca*) and European eel (*Anguilla anguilla*) with fin clips taken from all net-captured individuals. For glass eel (*A. anguilla*), individuals were sampled at coastal sluice gates by netting, and whole specimens were preserved. For mussel (*Mytilus edulis*), Pacific oyster (*Magallana gigas*) and flat oyster (*Ostrea edulis*), known populations of colonies were sampled, the shells of individuals were opened and soft tissue was collected. For Pacific oyster, which is not native to the Netherlands but was introduced at the tail end of the 1960s because of collapsing native flat oyster stocks, we sampled cultured populations from localities in the Waddenzee and Oosterschelde as well as ‘wild’ (i.e. unmanaged) populations. All samples were immediately preserved in 70% ethanol prior to DNA extraction in the laboratory and their GPS coordinates (decimal longitude and latitude) were recorded.

### DNA extraction, sequencing and bioinformatic processing

DNA was extracted from all samples using a DNeasy Blood and Tissue Kit following the manufacturers guidelines, with the final product eluted in 100ul TE buffer and stored at -80 °C. Extracted DNA was quantified using Qubit to ensure that it met a minimum molecular weight concentration of 10ng/ul, leaving a total of 2,152 individuals from 113 unique localities after excluding samples that failed to meet this criterion; turbot (*Scophthalmus maximus*, n=358 from 56 localities), zander (*Sander lucioperca*, n=152 from 2 localities), and European eel (*Anguilla anguilla*, n=606 from 17 localities), mussel (*Mytilus edulis*, n=550 from 11 localities), Pacific oyster (*Magallana gigas*, n=178 from 10 localities). SNP data for the flat oyster (*Ostrea edulis*) was generated for another study using a targeted SNP chip and bioinformatically processed data is re-used here (n=308 from 7 Dutch localities). For a full description of flat oyster sample collection and genotyping see Valk et al. (2026). Full details of species sampled including range and mean per locality are reported in Table 1. Full information including GPS points are reported in Table S1. For turbot, European eel, zander and mussel the samples were then sent to a commercial company (AgResearch, New Zealand, http://www.agresearch.co.nz) for GBS (Genotyping By Sequencing). For the flat oyster and the Pacific oyster, data was generated using a medium-density SNP array covering 14,950 SNPs for *O. edulis* and 40,625 SNPs for *M. gigas* (Gutierrez et al., 2017). Output genotypes (ped format) were exported using Axiom Analysis Suite (v.4.0.3, Thermo Fisher Scientific). For GBS samples, each was uniquely barcoded using Illumina adapters and multiplexed together in a GBS library. A total of ten GBS libraries were constructed (5 for Mussel, 2 for turbot, 2 for European eel and 1 for zander) according to the methods outlined in (Elshire et al., 2011) with modifications as outlined in (Dodds et al., 2015). Libraries were prepared using a *PstI*-*MspI* double-digest and included negative control samples (no DNA). Libraries underwent a Pippin Prep (SAGE Science, Beverly, Massachusetts, United States) to select fragments in the size range of 220-340 bp (genomic sequence plus 148 bp of adapters), and sequenced paired end on an Illumina NovaSeq 6000.

**Table 1.** Information on all samples genetically analysed in this study.

| Species | Sample type | Number of localities | Number of individuals | Range and mean of samples per population | Reference genome | Genome size (Gb) | Number of SNPs |
|---|---|---|---|---|---|---|---|
| turbot | Fin clip | 56 | 358 | 1-30 (mean=6.38) | GCA_963854745.1 | 0.56 | 9017 |
| European eel | Whole sample/ tail clip | 17 | 606 | 5-67 (mean=35.53) | GCA_013347855.1 | 1.36 | 66,521 |
| zander | Fin clip | 2 | 152 | 70-82 (mean=76) | GCA_023718345.1 | 1.14 | 2796 |
| Blue mussel | Muscle | 11 | 550 | 50 (mean=50) | GCA_963676885.2 | 1.7 | 3257 |
| flat oyster | Muscle | 7 | 308 | 31-62 (mean=44) | GCA_947568905.1 | 0.946 | 8403 |
| Pacific oyster | Muscle | 10 | 178 | 40-50 (mean=46.45) | GCF_963853765.1 | 0.648 | 30648 |

Raw reads in each GBS library were first trimmed to remove Illumina adapter sequences using Trimmomatic v.0.39 (Bolger et al., 2014) and then demultiplexed using the *process_radtags* module of Stacks v.2.53 (Rochette et al., 2019). Trimmed and demultiplexed reads were aligned to the species’ published (ENSEMBL) reference genomes using the *bwa-mem* algorithm (Li & Durbin, 2009) (GCA_963676885.2 for mussel, GCA_963854745.1 for turbot, GCA_013347855.1 for European eel, GCA_023718345.1 for zander) and processed in Stacks v2.53 (Rochette et al. 2019) to call SNPs. We ran a reference aligned analysis per species using the *ref_map*.*pl* perl script from Stacks, and retained only SNPs with a minor allele frequency of >0.05 that were present in at least 80% of sampled individuals per species, also removing loci that significantly deviated from Hardy-Weinberg Equilibrium (p<0.01) and only taking a single SNP per RAD locus to ensure that SNPs were not in strong linkage disequilibrium. For flat oyster and Pacific oyster, the genotyped reads were aligned with the reference genomes GCF_947568905.1 and GCF_963853765.1 respectively and followed the bioinformatic filtering procedures described in Valk et al. (2026). These SNP datasets were then analysed separately for each species and used for all subsequent analyses by converting to the required file formats (*.ped, *.map, *.structure, *.vcf) for each analysis. Because the datasets were generated using different genotyping approaches, analyses and interpretations were conducted primarily within species. Differences in marker ascertainment, genomic coverage and genotype quality can influence estimates of genetic diversity and relatedness; therefore, absolute values of these metrics were not interpreted as directly comparable across species or genotyping platforms.

### Population structure and genomic differentiation

To evaluate population structure, we performed a PCA in Plink v. 1.9.0 (Purcell et al., 2007) and plotted the samples on the first two principal components to visualise population clustering and variance explained by the PC axes. Additionally we ran Admixture v. 1.3.1 (Alexander et al., 2009) with *k* values ranging between 1 and 10 using a cross-validation approach and plotted ancestry coefficients per individual sample for each iteration of *k*. We calculated pairwise *F*_ST_ between sampling localities using the sample-size corrected approach of Weir and Cockerham (1984), implemented in Plink. For turbot, which included many localities where individuals were opportunistically sampled across the North sea, some with 1-2 individuals samples per locality, based on the PCA results we grouped localities for further analyses into *a priori* ‘Southern’, ‘Western’ and ‘Eastern’ clusters based on their geographic coordinates to simplify analysis and interpretation. Sampling localities for zander, European eel, mussel, flat oyster and Pacific oyster were treated as independent (i.e. not *a priori* grouped by geography).

### Genetic diversity, inbreeding and kinship

Individual-based observed (H_o_) and expected (H_e_) heterozygosity, nucleotide diversity (π) and inbreeding coefficients (F_is_) were calculated in Stacks v. 2.53 for all GBS datasets (turbot, blue mussel, European eel and zander). For flat oyster and Pacific oyster, H_o_, H_e_ and F_is_ were calculated using Plink, and nucleotide diversity (π) was calculated separately for each sampling locality using VCFtools v. 4.1 (Danecek et al. 2011) with the --window-pi 500000 option, retaining individuals from each locality using --keep. Kinship (pairwise relatedness, 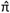) was calculated in Plink for all species and summarised by sampling locality. Allelic richness (A_r_) and private alleles (P_a_) were calculated using ADZE v. 1.0 (Szpiech et al., 2008), rarefying sample sizes to the lowest number of individuals per locality within each species, or within the a priori geographic groupings for turbot, to account for differences in sampling effort.

## Results

### Sequencing and bioinformatics

Over 4.03 billion sequence reads were obtained from the 10 GBS libraries across turbot, European eel, zander and mussel datasets. Of these, 2.84 billion reads were retained for further analysis after filtering out low quality reads and reads intersecting incomplete cut sites (1.34 billion for mussel, 1.06 billion for European eel, 276 million for turbot, 159 million for zander). Mean number of reads per sample per species (plus standard deviation) were 0.76 million (±0.32 million) for turbot, 1.73 million (±0.46 million) for European eel, 1.03 million (±0.28 million) for zander, 2.45 million (±0.59 million) for mussel. For Pacific oyster and flat oyster, following genotyping on the GENTYANE platform of INRAE Clermont-Ferrand, raw genotype calls were filtered for a minimal dish quality control (DQC) rate ≥ 0.78, and quality call (QC) rate ≥ 0.94, following best practices recommended by Affymetrix (Gutierrez et al., 2017). Final bi-allelic SNP datasets per species contained 9017 SNPs (turbot), 66521 SNPs (European eel), 2796 SNPs (zander), 3257 SNPs (mussel), 30,648 SNPS (Pacific oyster), and 8403 SNPs (flat oyster) after quality controls.

### Population structure and genomic differentiation

PCA and ADMIXTURE analyses revealed no strong evidence of population structure in the three fish species or Pacific oyster at the spatial scale examined (Fig. 2A–C, F). In contrast, mussel and flat oyster showed clearer genetic differentiation among sampling locations (Fig. 2D–E). These results indicate that genetic differentiation was weak or undetectable in the sampled fish populations and Pacific oyster, whereas more pronounced differentiation was evident in the two native and non-native bivalve systems examined. These results were supported by Admixture analyses (Fig. S1), and furthermore by estimates of pairwise *F*_ST_ (Fig. S2, Table S2), as well as by examining additional PC axes in the PCA (Fig. S3). Across taxa, these results indicate weak or absent population structure in fish species and Pacific oyster, contrasted with clear genetic differentiation among localities in mussel and flat oyster.

**Fig. 1.**
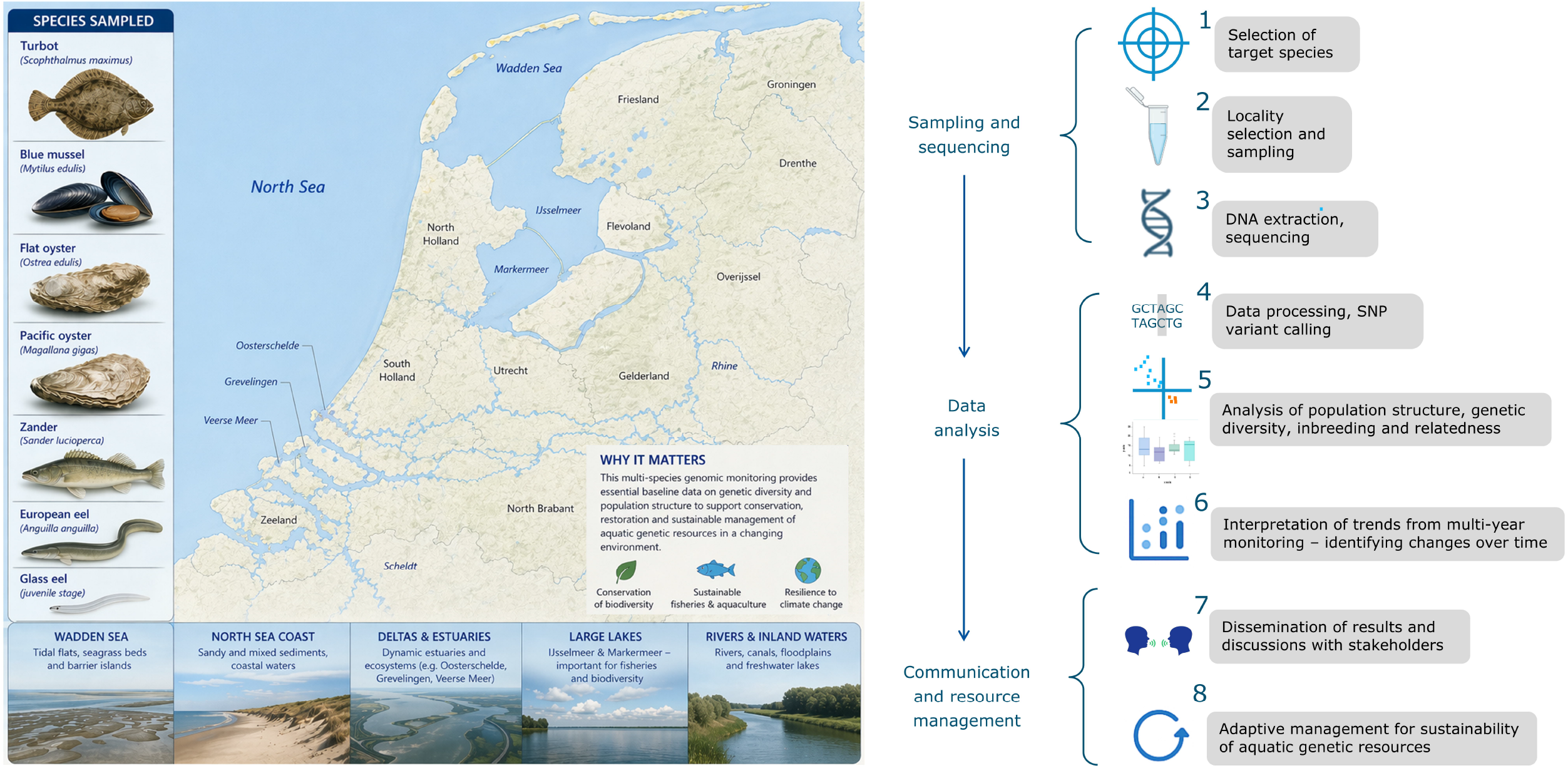
Overview of the Aquatic genetic resources (AqGR) management programme for the Netherlands, beginning with Sampling and sequencing, through Data analysis steps, ultimately to provide outputs for Communication and Resource management. Current completed aspects of the project presented in this manuscript encompass points 1 to 5, awaiting future time series data for monitoring temporal trends in population structure and genetic diversity which will guide points 6, 7 and 8.

**Fig. 2.**
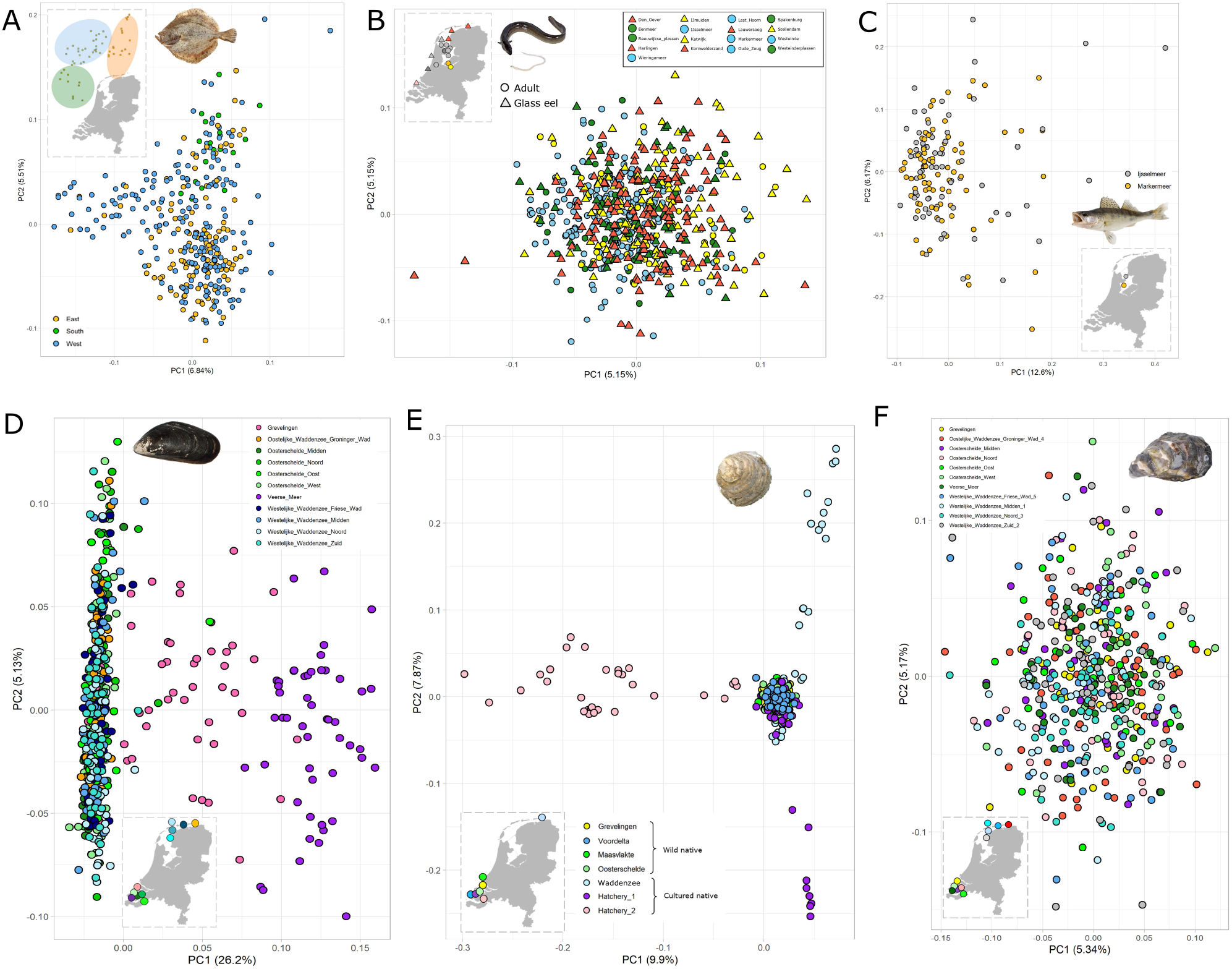
PCA based population structure analysis per species A) turbot, B) European eel, C) zander, D) mussel, E) flat oyster, F) Pacific oyster. Maps of sampling localities across the Netherlands are inset (dotted lines), coloured to match the PCA plots. Sampling locality names and types are provided per species for reference.

### Genetic diversity, inbreeding and kinship

For all three fish species, genetic diversity patterns were relatively homogeneous, consistent with the panmictic population structure detected. In turbot (Fig. 3A), minor differences in nucleotide diversity (π), private alleles (P_a_), and observed heterozygosity (H_o_) between localities were observed without a consistent geographic pattern. Similarly, in zander (Fig. 3C), differences were small, with slightly higher diversity in IJsselmeer. In European eel (Fig. 3B), coastal sampling localities (11–17), where glass eel were collected showed higher π and P_a_, with H_o_ also generally higher than in localities where adults were sampled (1–10). The differences remained when analyses were restricted to SNPs shared between the glass-eel and adult datasets. This sensitivity analysis suggests that the observed pattern is not explained solely by differences in the SNP sets analysed, although it cannot exclude all effects of library preparation, genotype quality or other sampling-related differences. In mussel, Grevelingen showed lower observed heterozygosity and higher estimated nucleotide diversity and private-allele counts than several other sampled locations (Fig. 3D). These are descriptive differences; their statistical significance was not formally evaluated. Flat oyster genetic diversity was generally homogeneous across localities, although Hatchery 2 showed a notably higher number of private alleles (Fig. 3E). Cultured populations overall exhibited higher relatedness and slightly lower nucleotide diversity compared to wild populations. For Pacific oyster, nucleotide diversity was similar across all localities, while private alleles were more abundant in northern (Waddenzee) sites, particularly Westelijke Waddenzee Zuid.

**Fig. 3.**
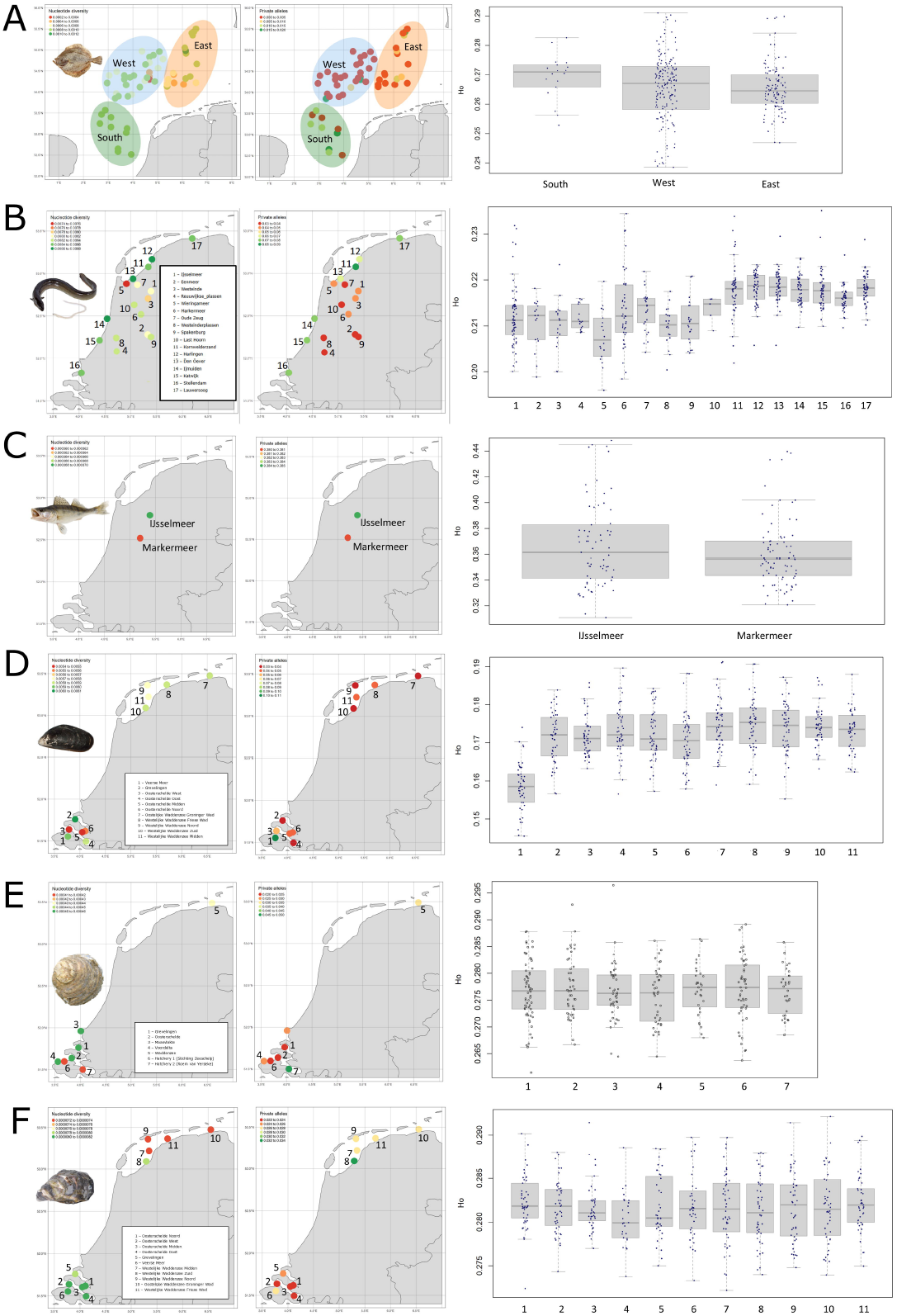
Genetic diversity metrics per species – maps of nucleotide diversity (π) per sampling locality (left), private alleles (P_a_) per sampling locality (centre), and boxplots of observed heterozygosity (H_o_) (right). For maps, colours of sampling localities represent the values (low-high, red-green). Each sampling locality is numbered/labelled on the maps corresponding to the boxplots, and scale of values for nucleotide diversity is shown in the top left of each map plot for A) turbot, B) European eel, C) zander, D) mussel, E) flat oyster, F) Pacific oyster.

Estimated F_is_ values were generally low in turbot, zander and flat oyster, and were somewhat higher in Pacific oyster (Fig. 4). These estimates indicate deviations from Hardy–Weinberg expectations at the sampled locations but should not, on their own, be interpreted as direct evidence of inbreeding, because population mixture, non-random sampling and genotyping-related effects can also contribute to heterozygote deficits. In European eel, F_is_ values were more variable and reached higher levels (F_is_ < 0.3), although coastal glass eel localities showed consistently lower values (F_is_ < 0.15) than inland adult sampling sites. In contrast, mussel exhibited substantially higher F_is_ values overall, with particularly elevated levels observed in Veerse Meer (F_is_ > 0.3). Pa erns of relatedness 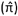 were broadly consistent with inbreeding estimates. For turbot, European eel, mussel, Pacific oyster and flat oyster, the majority of pairwise comparisons showed low relatedness 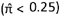, typically close to zero. However, several elevated 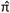 values were observed in European eel, suggestng the presence of potental sibling or half-sibling relatonships. In zander, mean 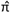 values were higher (approximately 0.25), consistent with a greater proportion of half-sibling relationships (Fig. 4A–F). All genetic diversity, inbreeding, and relatedness metrics per species and locality— including nucleotide diversity (π), private alleles (P_a_), observed heterozygosity (H_o_), inbreeding (F_is_), and relatedness 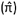 are provided in Table S3.

**Fig. 4.**
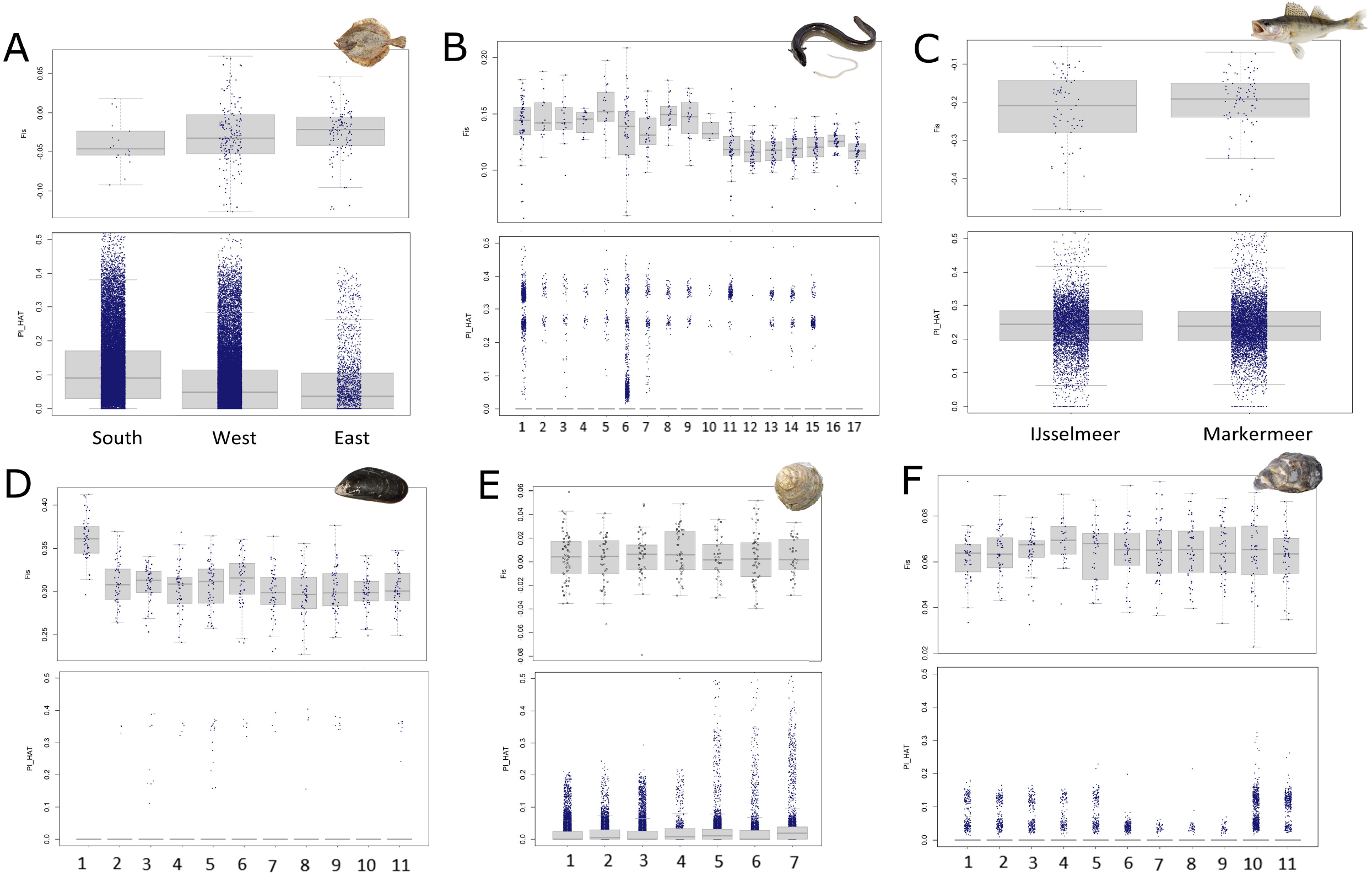
Inbreeding (F_is_, upper boxplot) and kinship (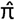, lower boxplot) metrics across sampling localities per species. Each data point for inbreeding represents an individual, each data point for kinship represents a pairwise comparison between individuals. A) turbot, B) European eel, C) zander, D) mussel, E) flat oyster, F) Pacific oyster.

## Discussion

Our results establish a genomic baseline for understanding and managing aquatic genetic resources applied at a national level, in this case, the Netherlands. Across the six fish and bivalve species we selected for their ecological and economic importance, we reveal both widespread connectivity and some unexpected local differentiation, as well as disparities in genetic diversity that may have direct implications for the management of these aquatic genetic resources. These findings underscore the value of multi-species genomic monitoring for identifying reservoirs of genetic variation and informing management decisions. In the sections that follow, we explore patterns of population structure and genetic diversity (including inbreeding and relatedness) across species, discuss their relevance for conservation strategies, consider potential biological and technical drivers of observed patterns, and highlight future directions for integrating genomics into long-term monitoring frameworks and resource conservation.

### Standing genetic diversity, inbreeding, relatedness and population structure across species

Patterns of genetic diversity and population structure differed markedly between fish and bivalves. In the sampled fish species, the lack of strong detectable population structure is consistent with substantial connectivity at the spatial scale examined. However, an absence of clear clustering does not demonstrate panmixia, and weak differentiation may also reflect the spatial scale of sampling, marker characteristics or limited representation of some localities. These results should therefore be interpreted as evidence of weak detectable structure rather than proof of unrestricted gene flow among all sampled locations. Such patterns are consistent with life-history traits including high dispersal capacity, large effective population sizes, and the absence of strong physical barriers to gene flow. These patterns align with previous knowledge for eel (Palm et al. 2009, Als et al. 2011, Enbody et al. 2021), turbot (Nielsen et al. 2004, Florin & Hoglund 2006) and zander (Eschbach et al. 2014, Béres et al. 2024) at similar spatial scales of sampling. Despite this, subtle variation in genetic diversity among sampling locations indicates that some areas may contribute disproportionately to overall diversity, highlighting the importance of considering local contributions even within broadly connected systems. In European eel (Fig. 3B), coastal sampling localities, where glass eel were collected, consistently showed higher π and P_a_, with H_O_ also generally higher than in localities where adults were sampled. Because glass eel and adult samples were processed in separate GBS libraries, we ensured that this pattern was unlikely to be due to technical artefacts (i.e. batch effects) by restricting analyses to utilise SNPs shared between datasets (see Paris et al. 2017, Crotti et al. 2020). These differences may reflect variation in the composition of the sampled groups, including differences in life stage, sampling environment or recruitment cohort, but the present sampling design does not allow these explanations to be distinguished. Dams and sluices can impede European eel migration in the Netherlands (Huisman et al. 2023), making connectivity an important consideration for eel management. However, our results do not establish whether migration barriers contribute to the differences in genetic diversity observed between coastal glass-eel and inland adult samples. In contrast, bivalve species (with the exception of the introduced Pacific oyster which shows no clear population structure) showed more pronounced population differentiation, consistent with more limited effective dispersal, local environmental gradients, and the influence of anthropogenic activities. For example, the strong differentiation observed in mussel populations may be associated with contrasting environmental conditions (e.g. salinity regimes, Westerbom et al. 2002) and varying degrees of hydrological isolation, which can restrict gene flow and promote local adaptation. Future sampling that separates life stage, location and sequencing batch, ideally with repeated sampling across sites and years, will help determine whether these differences reflect persistent spatial patterns, temporal variation or technical effects.

Alternatively, the difference in culture method (suspended rope culture vs. seabed bottom culture) could influence the genetic diversity between localities by affecting the dispersal and settlement of broadcast-spawned larvae, leading to differences in which individuals successfully recruit into each population (Demmer et al. 2022, Corrochano-Fraile et al. 2024). In flat oyster (*Ostrea edulis*), the distinction between cultured and wild populations, together with elevated relatedness in hatchery stocks, is consistent with reduced effective population sizes driven by founder effects, genetic drift, and non-random mating in hatchery systems (Lind et al. 2009; Vercaemer et al. 2006; Tonk et al. 2025). The presence of unique or shifted allele frequencies in cultured populations further suggests that hatchery practices can reduce genetic diversity and restructure population genetic composition, with potential consequences for fitness and adaptive capacity (Vercaemer et al. 2006; Tonk et al. 2025). The strength of inference differs among species because sampling coverage was uneven. In particular, the small number of individuals available from some turbot locations limits the resolution of location-specific comparisons, while the two sampled zander locations provide only a restricted assessment of spatial structure. Broader sampling will be needed to determine whether the observed patterns extend across the full geographic range of these species in the Netherlands.

### Management implications and conservation priorities

The observed patterns suggest contrasting management strategies across taxa. For species with weak population structure, such as turbot, European eel, zander and Pacific oyster, management may focus on maintaining overall genetic diversity at broader spatial scales. However, even in these systems, the uneven distribution of genetic diversity highlights the importance of protecting key source populations and maintaining connectivity to prevent the erosion of genetic variation over time. In highly connected species, management actions that disrupt migration routes or reduce population sizes could have system-wide genetic consequences, especially for example flood control measures such as sluices that disrupt flow conditions and may have negative effects on recruitment and survival (Huisman et al. 2023, Bozeman et al. 2024, Stoffers et al. 2022). For Pacific oyster in particular, which is an introduced non-native species in the Netherlands, the lack of clear population structure is likely linked to its recent introduction and capacity to rapidly adapt to new environments. For bivalves, stronger population structuring indicates that conservation efforts should prioritise the maintenance of multiple, distinct populations rather than treating them as a single management unit. This is particularly important where populations may be locally adapted to specific environmental conditions. The mussel population in Veerse Meer illustrates a potential trade-off between preserving unique, possibly locally adapted variation and mitigating inbreeding through increased connectivity. Such trade-offs are central to conservation genetics, where interventions aimed at increasing gene flow must be balanced against the risk of outbreeding depression or loss of local adaptation. In flat oysters, the differentiation exhibited by cultured populations in hatcheries and in the Waddenzee likely reflects a combination of founder effects, drift, and hatchery practices. These findings highlight the need for careful broodstock management and genetic monitoring in restoration and aquaculture programmes to avoid unintended reductions in genetic diversity or increases in relatedness. Incorporating genetic criteria into breeding and translocation strategies will be essential to ensure long-term population viability.

### Towards long-term monitoring of aquatic genetic resources

This study establishes a foundation for the long-term monitoring of aquatic genetic resources (AqGR) in the Netherlands, providing baseline estimates of genetic diversity, inbreeding, relatedness, and population structure across multiple species of ecological and fisheries relevance. Such baselines are critical for the sustainable management of aquatic biodiversity, as genetic diversity underpins adaptive potential, population viability, and long-term productivity of both wild and cultured stocks (Allendorf et al. 2013; Hoban et al. 2020). In this context, genomic monitoring is increasingly recognised as a key component of ecosystem-based fisheries management and aquaculture sustainability, directly linking biodiversity conservation with food security and blue economy objectives (FAO 2019; Bernatchez et al. 2017).

The multi-species framework adopted here is scalable and transferable to other regions and taxa, providing a model for integrating genomics into national and international AqGR monitoring programmes. This aligns with global calls to operationalise genetic indicators within biodiversity reporting frameworks and to ensure sustainable use of aquatic genetic resources in support of resilient food systems (FAO 2019; Hoban et al. 2022). Continued monitoring will enable detection of temporal changes in genetic diversity, including risks such as inbreeding, loss of adaptive variation, and increased relatedness, which can reduce population productivity and adaptive capacity in both wild fisheries and aquaculture broodstocks (Laikre et al. 2010; Hare et al. 2011).

These dynamics are particularly relevant in aquatic systems where environmental change, altered connectivity, and harvest pressure can rapidly reshape population structure and recruitment dynamics, with direct implications for fisheries yields and aquaculture performance (Selkoe & Toonen 2011; Bernatchez et al. 2017). While this study provides a robust genomic baseline, limitations remain. For zander, sampling was restricted to only two localities, which limits the ability to resolve fine-scale spatial genetic structure and to distinguish discrete population structuring. This restricted design also reduces power to detect subtle patterns of connectivity and gene flow among local spawning groups and precludes assessment of temporal stability in allele frequencies and genetic diversity. Expanding both spatial and temporal coverage will therefore be essential for accurately characterizing population structure, identifying biologically meaningful management units, and detecting trends in connectivity and exploitation relevant to fisheries management. In addition, reduced-representation sequencing approaches such as GBS and SNP chips enabled efficient multi-species genotyping at scale but present challenges for long-term standardisation, particularly due to locus dropout, batch effects, and variation in marker representation across libraries (Davey et al. 2011; Lowry et al. 2017). These issues may affect temporal comparability in long-term AqGR monitoring programmes if not carefully controlled. Future efforts should increasingly incorporate whole-genome sequencing approaches, which provide more complete genomic representation and enable more accurate estimation of inbreeding (e.g. via runs of homozygosity), genetic load, and adaptive variation (Kardos et al. 2015; Ceballos et al. 2018). Such approaches are particularly valuable for fisheries and aquaculture systems, where maintaining genetic diversity is directly linked to stock resilience, production stability, and food security under global change (FAO 2019; Bernatchez et al. 2017).

## Acknowledgements

This research was carried out by the Centre for Genetic Resources the Netherlands (CGN) and funded by the Ministry of Agriculture, Fisheries, Food Security and Nature (LVVN), within the framework of WOT-03 Genetic Resources (project number WOT-03-002-068 and WOT-03-001-069). We thank Edward Schram at Wageningen Marine Research (IJmuiden) and Ainhoa Blanco (Yerseke), as well as Waddenunit and Rijksrederij (Luctor and Regulus) for assistance obtaining samples.

## Data accessibility and code

All raw genomic data is uploaded to ENA as PRJNAxxxxxxx. A DRYAD repository containing raw processed data and a linked ZENODO repository containing all code needed to generate the results obtained in this manuscript can be found here: xxxxxxx

