## Supplementary figures and images for "A genomic baseline for long-term monitoring of economically important aquatic species in the Netherlands"

### Fig S1

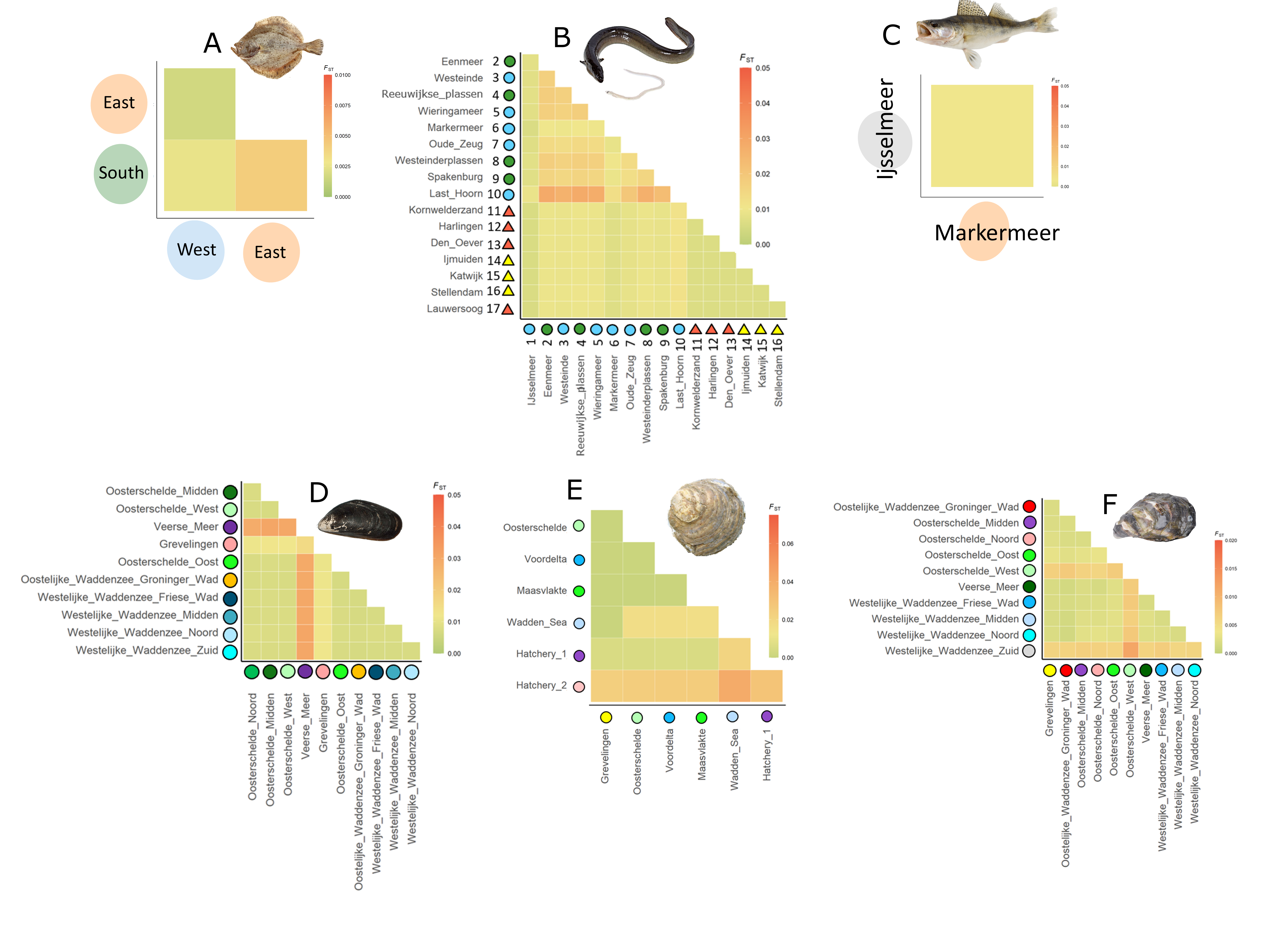

### Fig S2

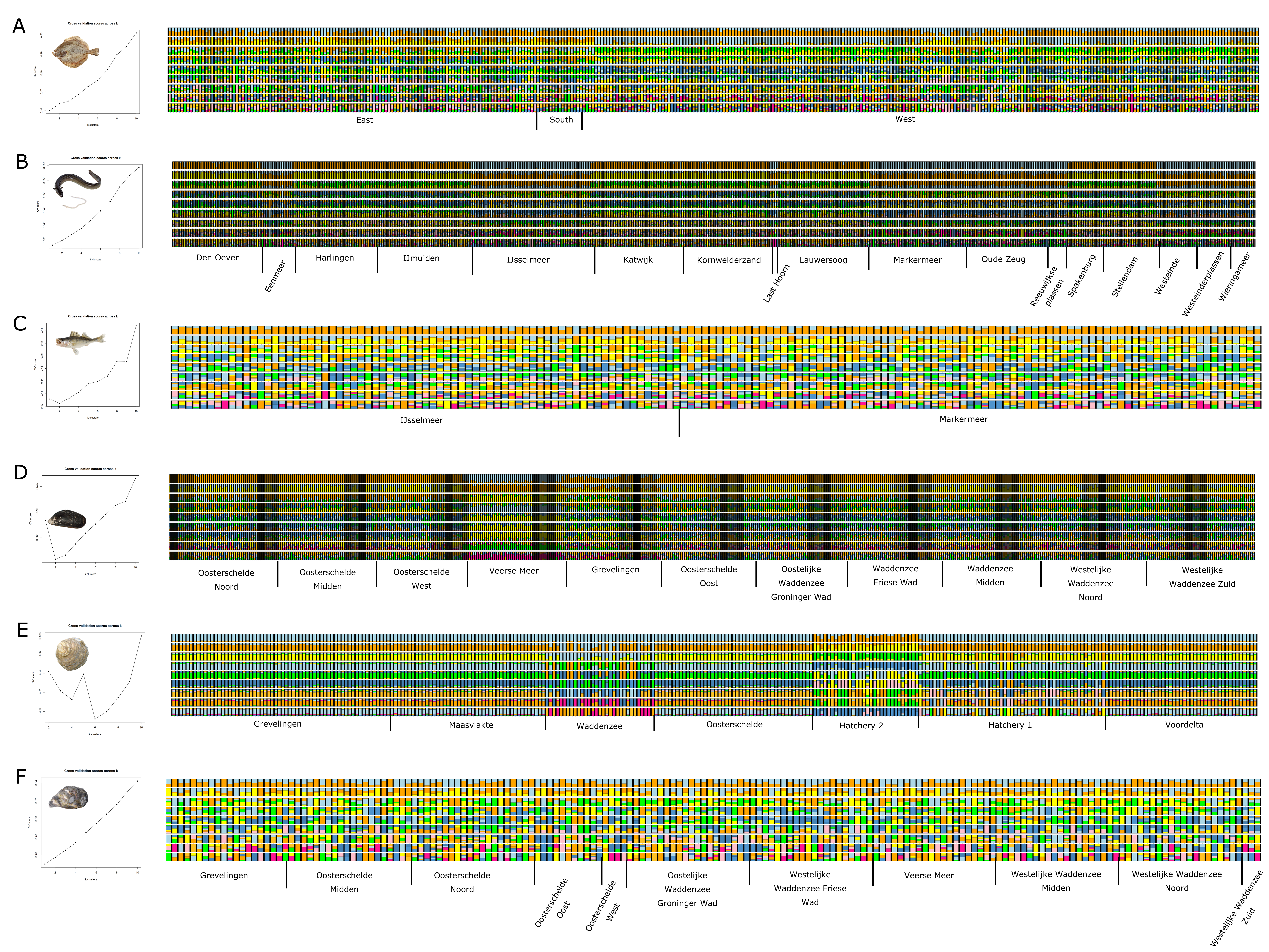

### Fig S3

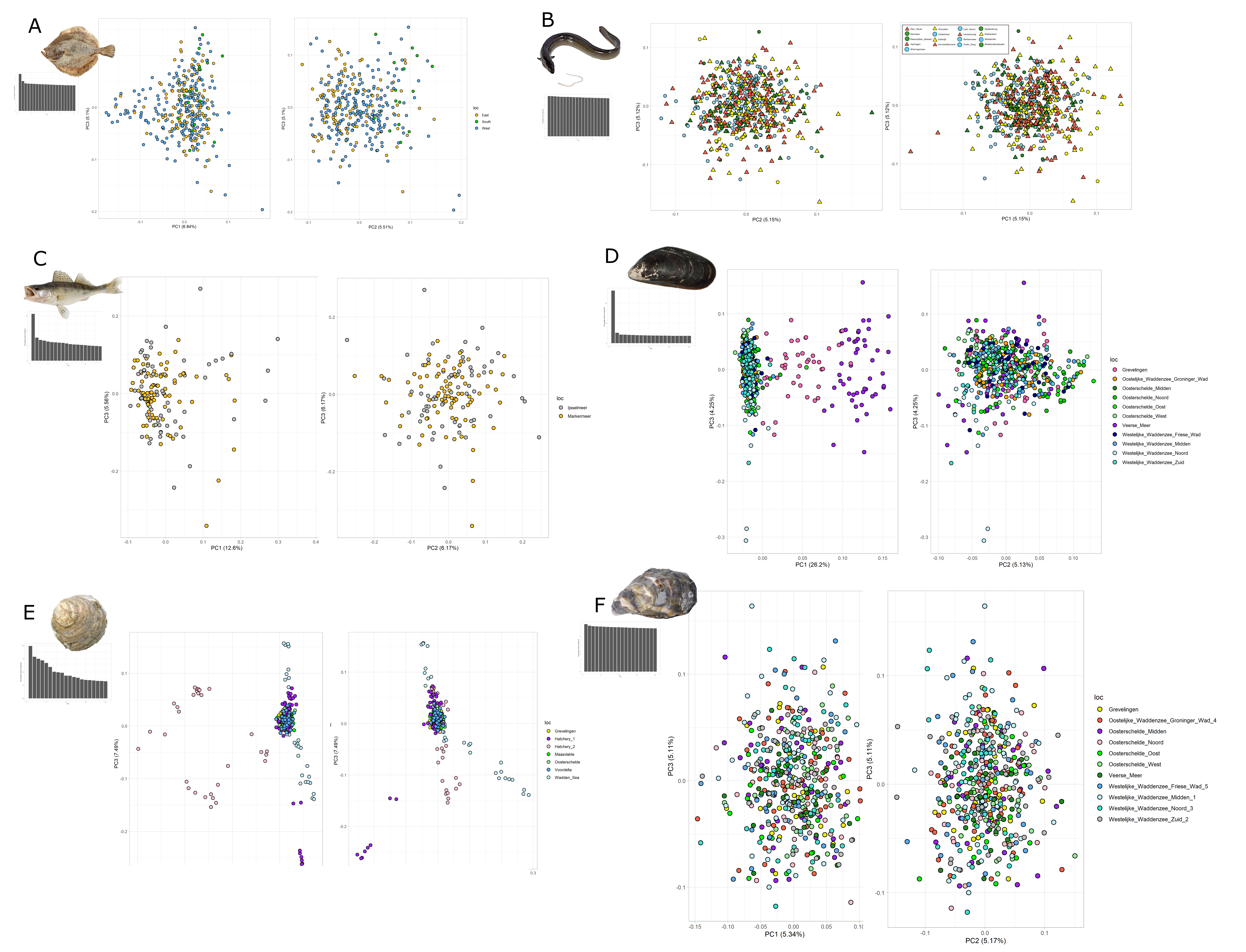
